# Accurate haplotype-resolved *de novo* assembly of human genomes with RFhap

**DOI:** 10.64898/2026.01.28.702238

**Authors:** Damariz Gonzalez, Gabriel Cabas, Juan Francisco Miquel, Carol Moraga, Francisca Salas, Alex Di Genova

## Abstract

Haplotype-resolved *de novo* assemblies enable genome-wide separation of maternal and paternal variation, improving the interpretation of complex variants relevant to human disease. Trio-aware assemblers such as Hifiasm-Trio leverage parental short reads by deriving parent-specific k-mers to guide phasing within the long-read assembly graph; however, fixed k-mer length and heuristics cannot be optimal in the presence of sequencing errors and graph complexity in repetitive regions, contributing to phasing errors and reduced haplotype-resolved contiguity. Here we present RFhap, a trio-based long-read phasing method that integrates multi-k-mer parent-specific markers with an alignment-free k-mer lookup engine and a random forest classifier to assign long reads to maternal, paternal, or unknown haplotypes prior to *de novo* assembly. We benchmarked RFhap on four human trio datasets from the Human Pangenome project spanning two ONT chemistries (R10.4 and R9.4.1). Using Merqury to evaluate downstream assemblies, RFhap nearly doubled corrected haplotype NG50 (mean 24.3 Mb vs 13.1 Mb) and halved switch error rates (mean 0.111% vs 0.236%) relative to Hifiasm-Trio, while remaining competitive in consensus QV and parental k-mer completeness. Consistent with improved phasing, RFhap reduced long-switch errors by ∼3-fold across datasets, particularly within interspersed repeats. Together, these results demonstrate that RFhap improves phasing accuracy from standard trio data, representing a step toward accurate and automated diploid human *de novo* assembly from long reads.

## Introduction

Advances in current sequencing technologies, specifically long-read sequencing, have enabled the construction of more continuous *de novo* assemblies, capitalized by the first complete telomere-to-telomere human genome[1]. Long-read technologies such as Oxford Nanopore [2] and Pacific Bioscience [3], generate long-range information that has allowed better haplotype-resolved assemblies [4][5][6] and a higher resolution of genetic variation associated with disease[7,8][4]. However, current long-read haplotype-resolved *de novo* assembly methods, such as Hifiasm [5] and Verkko [6], still need to improve to achieve automated diploid *de novo* assembly from long reads [9,10]. Although phasing blocks at the scale of megabase are routinary achieved[5][6], switch phase errors, which are hard to error-correct[11], as well as the need for multiple sequencing technologies[4,11][5][6] and computational resources[5,11][6], are still open problems to routinary obtain accurate haplotype-resolved chromosome-scale diploid assembly of human genomes.

The recently completed T2T-HG002 diploid genome[11] illustrates these limitations and the extent of current technical requirements. This assembly integrated ultra-high long-read coverage data[12,13] (∼170× PacBio HiFi and ∼209× ONT ultra-long reads) with Illumina trio data from both parents, as well as Strand-seq[14] and Hi-C[15] data to enable accurate phasing and scaffolding across complex genomic regions. An initial Verkko[6] assembly was iteratively refined by closing gaps with reference genomes[1], resolving complex regions via manual graph inspection[11], and assembling specialized loci such as rDNA with dedicated tools[16]. Three rounds of polishing[17,18], guided by multiple independent sequencing technologies[14,15] and crowdsourced curation[11], corrected tens of thousands of base-level, small indel errors and introduced larger consensus patches while preserving global structural accuracy[11]. This workflow demonstrates that truly complete, diploid, telomere-to-telomere references currently require multiple complementary sequencing technologies, reference genomes, intensive error correction, and manual curation, as well as substantial computational resources [11].

Current *de novo* assemblers can integrate external sources of haplotype information, most commonly parental (trio) sequencing data [19], Strand-seq [14], and Hi-C [15]. Among these options, trio-based approaches are experimentally the most straightforward, as they rely on standard short-read sequencing of both parents and the offspring[19], without the need for additional specialized protocols. In early trio-based strategies, such as trio-binning [19], parental short reads are used to derive parent-specific k-mers that partition offspring long reads by parental origin before assembly, effectively transforming a diploid genome into two separate haploid assembly partitions. Hifiasm[5] and Verkko[6] use trio data more directly within the assembly graph; the parental k-mer information is used to decorate the graph, enabling the phasing of heterozygous sites and the handling of switch errors spanning megabase-long contigs. However, trio-based methods typically rely on a single, fixed k-mer length (often k = 21) to define parental markers[5,6,19], which may be suboptimal in terms of both sensitivity and robustness to sequencing errors. Moreover, assembly graphs remain highly tangled in repetitive regions [5,6], and decorating such graphs with sparse or parental markers can be error-prone. In practice, haplotypes are often assigned by comparing the counts of paternal versus maternal markers on reads[19] or graph edges[5,6]; when marker counts are low, imbalanced, or affected by errors, this majority-vote scheme can misclassify haplotypes. Collectively, these heuristics contribute to increased switch error rates and suboptimal phasing of *de novo* diploid genome assemblies.

Here we introduce RFhap, a trio-based phasing method that combines multi–k-mer information with machine learning to improve long-read haplotype assignment. RFhap uses standard trio data and an alignment-free k-mer engine to look up parental-specific k-mers of multiple lengths in offspring long reads. These multi–k-mer hits and counts are then integrated in a Random Forest classifier that assigns each long read to the maternal, paternal, or unknown haplotype. Haplotyped reads are assembled by partition using hifiasm. Benchmark of RFhap on four human datasets evaluated with Merqury[20] using an automated Nextflow pipeline demonstrates that RFhap nearly doubled corrected haplotype NG50 (mean 24.3 Mb vs 13.1 Mb) and halved switch error rates (mean 0.111% vs 0.236%) relative to Hifiasm-Trio, while remaining competitive in consensus QV and parental k-mer completeness, representing a step toward accurate and fully automated diploid de novo assembly from long reads and trio data. The RFhap pipeline is implemented in NextFlow for easy execution and reproducibility.

## Results

### The RFhap algorithm

RFhap classifies long reads from trio-based data using a five-stage process that couples parent-specific k-mer discovery with a machine-learning haplotype-aware read classifier and partitioning (See Methods). In brief, parental short reads are used to derive maternal- and paternal-exclusive k-mer sets (KMC3[21]), which are then queried along child ONT reads (FASTKM) to generate per-read marker statistics across multiple k values. These features train a multiclass random forest that assigns reads to either haplotype A, haplotype B, or an unknown category. The resulting labels are then used to partition reads for downstream haplotype-resolved assembly (Fig. S1).

The pipeline is compute-dominated by FASTKM (k-mer lookup), while kmc3 (k-mer set construction) contributes a smaller but I/O-heavy fraction; random-forest training/prediction and read partition are comparatively lightweight (Fig. S2). Overall, the classification process of RFhap completed in < 850 CPU-hours (≈6.3 hours elapsed) on the HG002 human whole-genome trio-dataset, with a peak memory requirement of ∼30 GiB needed by the k-mer/lookup steps (Fig. S2). This efficient runtime is enabled by implementing an alignment-free algorithm (k-mer membership lookup instead of read mapping) together with lightweight data structures (rolling hash[22], minimal perfect hash[23], and compact fingerprints[24] for constant-time k-mer queries), allowing RFhap to scale to human WGS trio-data while keeping compute and memory demands modest.

### RFhap is robust to sequencing errors

To evaluate RFhap, we analyzed four human trio datasets (HG002, HG005, HG01109, and HG01243) using parental short reads and child ONT long reads obtained from the Human Pangenome project [13] (Methods). Long-read data ranged from ∼160–294 Gb per child, comprising 7.6–16.3 million reads per sample, with mean read lengths of ∼12.5–24.3 kb and read N50 values spanning ∼23.9–52.9 kb (Sup. Table 1 and Fig. S3). The datasets included both R10.4 (HG002, HG005) and R9.4.1 (HG01109, HG01243) ONT chemistries, with average qualities of Q19 and Q15, respectively, enabling the assessment of RFhap across distinct error profiles and sequencing conditions.

Across the four trios, RFhap achieved consistently high testing model classification performance, with F1 scores ∼0.99–1.00 for all classes (A, B and U) and only a small variation between samples and ONT chemistries (Fig. 1A, Fig. S4). Performance was stable across the two sequencing chemistries (R10.4 vs R9.4.1), indicating that the model generalizes well across pore/kit versions and across individuals (Figure 1A). Feature-importance profiles were consistent across chemistries, but the model clearly reweighted which signals it trusted most (Fig. 1B–C). Under R10.4 (Q19), which has a lower raw read error rate than R9.4.1 (Q14), the classifier placed relatively more weight on a subset of high-discrimination k-mer/summary features (e.g., k33_A, totA, k33_B, k33_A_mean; positive shifts in Fig. 1C). In contrast, under R9.4.1, importance shifted toward features that appear more error-tolerant/aggregative in this setting (e.g., abk30/abk27, abkT, totB, k30_B; negative shifts in Fig. 1C), consistent with the model compensating for prone-error reads by relying more on robust, read-level or abundance-like signals rather than the same high-weight discriminators used in R10.4. Agreement analysis of read haplotype classification between RFhap, Hifiasm, and trio-binning shows that most read haplotype assignments are concordant across all three approaches: 411,000 reads (79.57%) fall in the three-way intersection (Fig. 1D). Among discordant cases, the largest subset corresponds to RFhap + trio binning agreement (62,290 reads; 12.06%), followed by trio binning + Hifiasm agreement (22,699; 4.39%), and a smaller set where RFhap + Hifiasm agree (8,498; 1.65%). Moreover, 12,039 reads (2.33%) were discordant between all methods. The impact of these method-specific read-haplotype assignments is determined by evaluating their effects on the continuity and accuracy of downstream haplotype-resolved *de novo* assembly.

**Figure 1.**
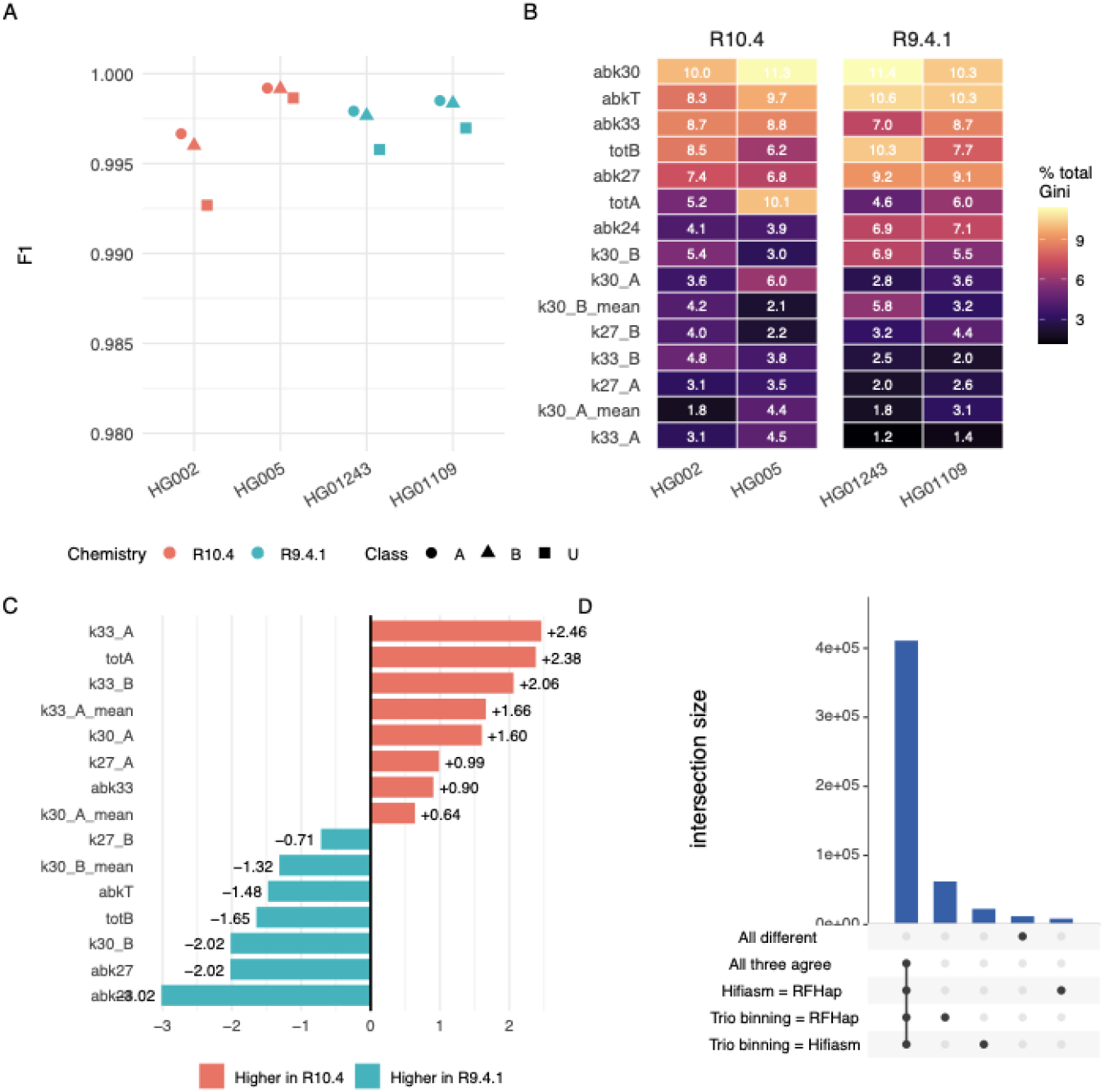
RFhap performance, feature importance, and agreement across ONT chemistries and samples. (A) F1 scores for the three classes (A, B, U; indicated by point shape) across four human genomes (HG002, HG005, HG01243, HG01109), stratified by ONT chemistry (R10.4 in red; R9.4.1 in teal). (B) Heatmap summarizing feature importance for each dataset, reported as the percentage contribution to the total Gini importance (values shown in each cell), grouped by chemistry (R10.4: HG002/HG005; R9.4.1: HG01243/HG01109). (C) Differences in feature importance between chemistries (R10.4 - R9.4.1), where positive bars (red) indicate features with higher importance under R10.4 and negative bars (teal) indicate features with higher importance under R9.4.1. (D) UpSet plot showing concordance patterns among RFhap, Hifiasm, and trio-binning haplotype assignments; bars denote intersection sizes for complete agreement, pairwise agreement, and cases where all three methods differ.

### RFhap improves phasing accuracy of haplotype-resolved de novo assembly

To assess the impact of RFhap read-haplotype assignments on *de novo* diploid assembly, we assembled four human trio datasets: HG002 and HG005 generated with Oxford Nanopore R10.4 chemistry, and HG01109 and HG01243 generated with Oxford Nanopore R9.4.1 chemistry. RFhap-partitioned reads were assembled with Hifiasm[5] (-ont option) and benchmarked against Hifiasm-Trio, using the same trio datasets for both methods. The resulting haplotype-resolved assemblies were evaluated with Merqury[20] to quantify switch error rates, corrected haplotype NG50, consensus quality (QV), parental k-mer completeness, and the type and distribution of phasing errors.

Across the four trio datasets, RFhap-based assemblies consistently improved haplotype contiguity and phasing accuracy relative to state-of-the-art Hifiasm-trio. The Merqury-corrected haplotype NG50 increased substantially for HG002 (35.4 Mb with RFhap vs 12.0 Mb with Hifiasm) and for the R9.4.1 datasets HG01109/HG01243 (∼14.8–15.6 Mb with RFhap vs ∼4.6–5.2 Mb with Hifiasm), while remaining comparable in HG005 (∼31.2 Mb RFhap vs ∼30.6 Mb Hifiasm, Figure 2A). Raw assembly contiguity (NG50) was overall comparable between methods(Figure 2A), with a modest improvement under RFhap across datasets (mean ∼76 Mb RFhap vs ∼73 Mb Hifiasm). Importantly, switch error rates were reduced in all datasets (Figure 2B), including HG002 (0.056% RFhap vs 0.094% Hifiasm), HG005 (0.047% vs 0.073%), and HG01109/HG01243 (∼0.165–0.175% vs ∼0.36–0.42%). Notably, the greatest improvements in corrected NG50 and switch error were observed for the ONT R9.4.1 assemblies compared to R10.4 (Figure 2), which is expected given that RFhap’s multi-k-mer strategy is designed to be more robust to sequencing errors during haplotype assignment.

**Figure 2.**
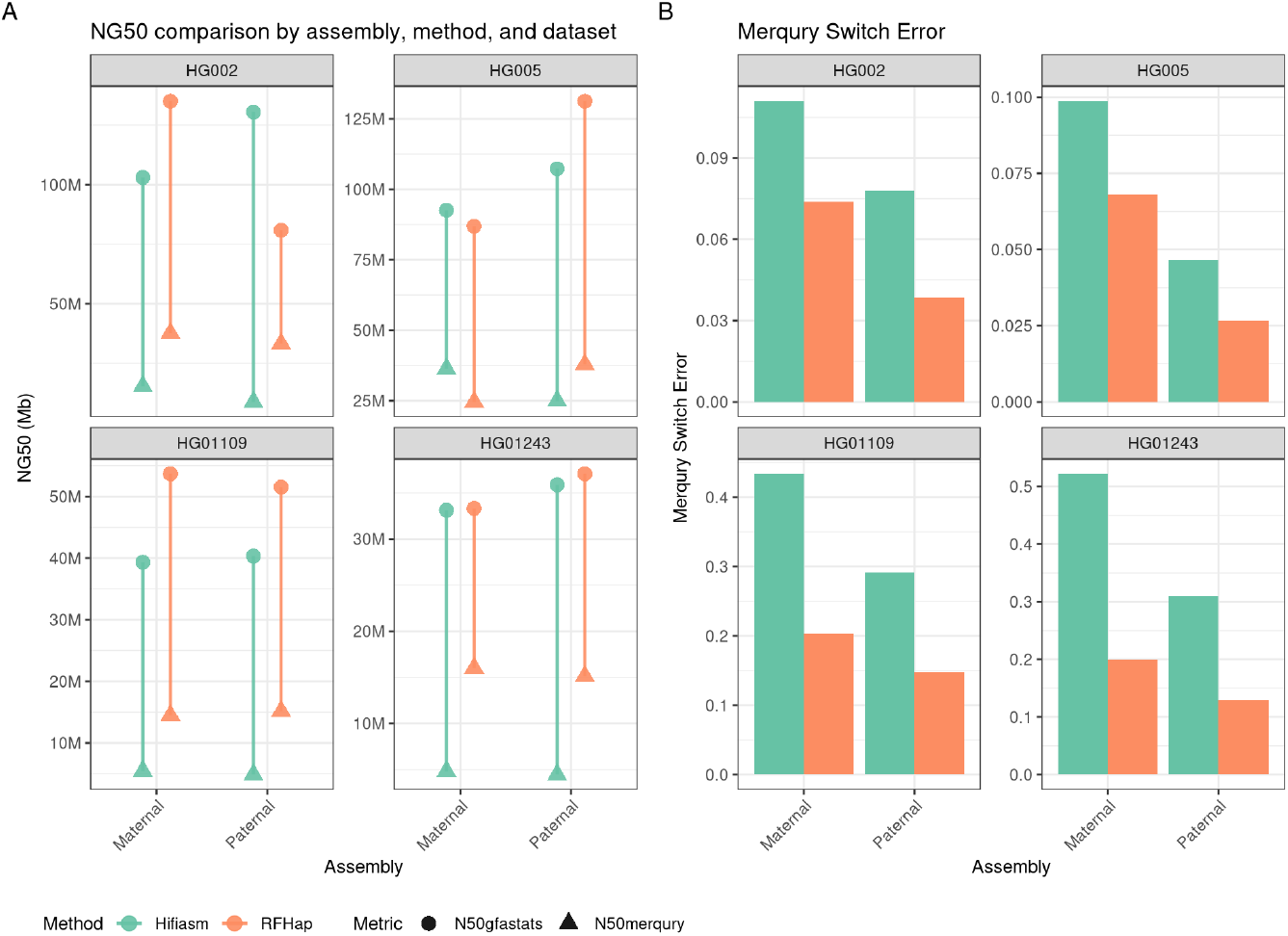
Comparison of assembly contiguity and phasing accuracy across datasets and methods. (A) N50 values for maternal and paternal assemblies across four datasets (HG002, HG005, HG01109, HG01243). NG50 is shown for two evaluation metrics: gfastats (circles) and Mercury corrected N50 (triangles). Points are colored by classification method(RFhap and Hifiasm trio mode). (B) Merqury Switch Error Rates for maternal and paternal assemblies across the same datasets, shown separately for each method. Lower values indicate improved phasing accuracy.

Assembly consensus quality (Merqury QV) was primarily driven by ONT chemistry (Figure S5A), with R10.4 datasets (HG002/HG005) achieving substantially higher QV than R9.4.1 datasets (HG01109/HG01243). Across R10.4 assemblies, Hifiasm showed the highest consensus accuracy (mean QV ≈ 57.5), whereas RFhap assemblies were slightly lower (mean QV ≈ 53.4). In contrast, for R9.4.1 assemblies, both methods produced similar QV values (Hifiasm mean QV ≈ 38.4 vs RFhap mean QV ≈ 37.6), indicating that the consensus quality gap between methods is modest under R9.4.1 chemistry. This pattern is expected, since RFhap partition reads before assembly, reducing the number of long reads available for consensus polishing, whereas Hifiasm-Trio use the full long-read set during consensus generation. Parental k-mer completeness mirrored the QV trends and was also chemistry-dependent (Figure S5B). For the R10.4 datasets (HG002/HG005), Hifiasm achieved higher overall completeness (∼98.2% mean) than RFhap (∼95.7% mean), consistent with its slightly higher consensus QV when using the full long-read set for polishing. In contrast, for R9.4.1 datasets (HG01109/HG01243), completeness was high and comparable between methods (∼97.0% Hifiasm vs ∼96.4% RFhap), matching the smaller QV gap observed under R9.4.1 chemistry.

Overall, we demonstrated that RFhap nearly doubled corrected NG50 (mean 24.3 Mb vs 13.1 Mb) while halving switch errors (mean 0.111% vs 0.236%) and is competitive in QV and k-mer completeness.

### RFhap reduces long-switch errors within interspersed repeats

The improved phasing accuracy of RFhap translated into markedly fewer Merqury long-switch errors (sustained switch blocks, see methods), with totals decreasing from 1,724 to 513 in HG002, 3,359 to 1,244 in HG01109, and 3,677 to 963 in HG01243; HG005 remained comparable (490 vs 467). RFhap achieved a ∼3× reduction in long-switch errors across the four trios (9,250 vs 3,187, Figure S6A). Merqury short errors also decreased, with an overall ∼2.7× reduction across datasets (Figure S6B).

To characterize the genomic distribution of these long-switch error regions, we extracted the flanking sequences of each long-switch error interval and mapped them to the CHM13 v2.0 T2T human reference[1]. Long-switch error regions were strongly enriched in interspersed repeats (SINE/LINE/LTR), which dominated the Hifiasm diploid assemblies (Figure 3). RFhap substantially reduced long-switch errors in these classes (Figure 3A), for example in HG002 (SINE 1381 to 412; LINE 1235 to 417; LTR 694 to 179), HG01109 (LINE 2468 to 954; SINE 2403 to 1127; LTR 1170 to 574), and HG01243 (LINE 2644 to 697; SINE 2628 to 785; LTR 1348 to 442). In contrast, satellite-associated long-switch errors were moderately more abundant in RFhap (and increased in some cases; e.g., HG01243 449 to 742, HG01109 610 to 680), indicating that after suppressing long-switch errors in interspersed repeats, the residual long-switch errors are increasingly concentrated in tandem/satellite-rich regions, which remain challenging to phase in human diploid assembly.

**Figure 3.**
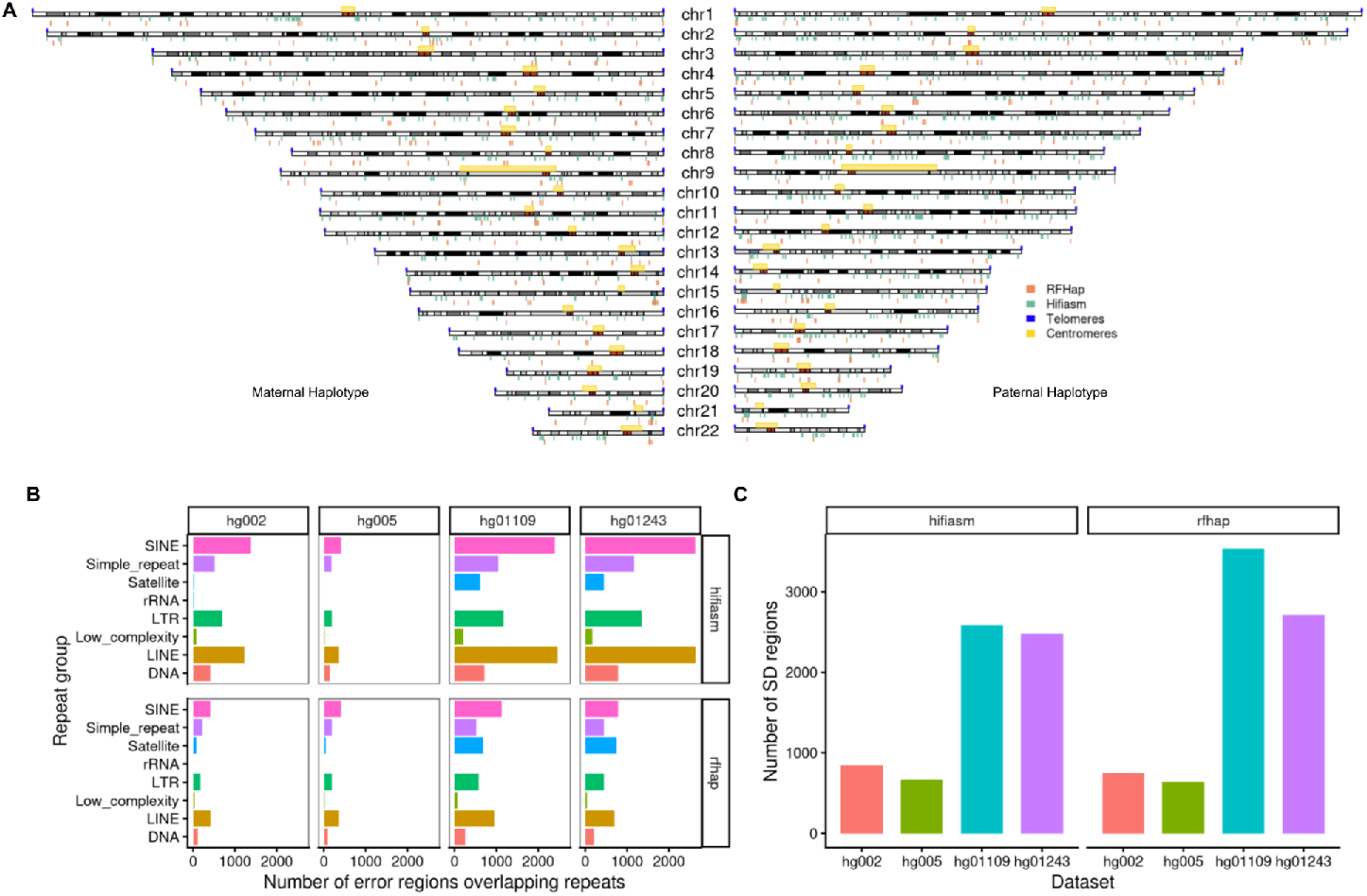
Genomic distribution of Merqury long switch-errors across repeat classes. (A) Chromosome-level maps of long switch-error regions for the maternal (left) and paternal (right) haplotypes. Error blocks are shown along each chromosome for RFhap (orange) and Hifiasm-trio (green), with telomeres (blue) and centromeres (yellow) indicated for reference. (B) Overlap of long switch-error regions with major repeat groups (DNA transposons, LINEs, SINEs, LTRs, low-complexity, simple repeats, satellites, rRNA) across the four trios (HG002, HG005, HG01109, HG01243), shown separately for Hifiasm-trio (top row) and RFhap (bottom row). (C) Number of long switch-error regions intersecting segmental duplications (SDs) per dataset for each method, highlighting the concentration of long-range phasing inconsistencies in duplication-rich regions.

Long-switch errors were also frequent in segmental duplications (SDs) (Figure 3C). RFhap slightly reduced SD-associated long errors in the R10.4 datasets (HG002 842 to 749; HG005 666 to 638), but SD overlaps increased in the R9.4.1 datasets (HG01109 2584 to 3531; HG01243 2476 to 2715). This is expected because SDs are highly similar, and the lower per-read accuracy of R9.4.1 makes haplotype phasing particularly challenging in duplicated regions.

In summary, RFhap strongly reduces long-switch errors genome-wide, with the largest improvements in interspersed repeats. The remaining long-switch errors are preferentially retained in tandem repeats and segmental duplications.

## Discussion

Accurate, fully automated diploid *de novo* assembly from long reads remains limited not by contig continuity per se, but by haplotype correctness. Modern assemblers such as Hifiasm and Verkko routinely generate megabase-scale phased blocks, yet switch errors persist and are difficult to correct. The T2T-HG002 diploid effort emphasizes this gap: achieving a complete, highly accurate diploid reference required high sequencing depth, multiple technologies, iterative polishing, and extensive manual curation. In this context, RFhap targets a core upstream problem: long-read-level haplotype assignment under sparse, error-prone k-mer signals, aiming to improve diploid assembly phasing using only standard trio data.

RFhap’s main implementation choice is to replace fixed-lengh parental-specific k-mers and majority-vote heuristics with a multi–k-mer feature representation integrated by a random forest classifier. Across four trios spanning two ONT chemistries, the classifier maintained excellent performance and showed chemistry-specific feature reweighting, consistent with an adaptive use of robust features when long-read error rates increase. This feature reweighting explains why downstream benefits were higher for R9.4.1 datasets, where sequencing errors can erode single-k sensitivity and inflate misclassification. Importantly, most long-read assignments were concordant across RFhap and Hifiasm, yet the relatively small discordant fraction still had measurable consequences at the assembly scale, supporting the idea that systematic misassignments can drive long-range phasing errors.

At the assembly level, RFhap improved haplotype-resolved quality primarily by reducing phasing errors rather than by altering raw contiguity. Across datasets, RFhap nearly doubled corrected NG50 and halved switch error rates, while raw NG50 remained similar. This separation between raw and “corrected” contiguity indicates that RFhap is not simply generating longer contigs, but producing contigs that remain consistent under haplotype-aware evaluation. Consensus quality metrics were dominated by ONT chemistry and, for R10.4, Hifiasm-trio retained a modest advantage. This is expected because RFhap partitions reads before assembly, effectively reducing depth for consensus generation notably in regions where the two haplotypes are identical, while Hifiasm-trio use the full read set during consensus generation. Notably, RFhap remained competitive in QV and completeness (especially under R9.4.1), suggesting that improved haplotype purity may partially offset reduced depth by limiting cross-haplotype mixing and associated polishing conflicts. Practically, these results imply that RFhap can improve phasing without imposing a major penalty on consensus accuracy, but that consensus polishing strategies tailored to partitioned read sets can potentially close the remaining gap under high-quality chemistries.

Long-switch errors provide a mechanistic view of where diploid phasing breaks down. RFhap achieved an overall ∼3× reduction in Merqury long-switch errors, and reduced short errors as well. Mapping long-switch intervals to CHM13 v2.0 showed that, in Hifiasm-trio assemblies, long-switch errors were enriched in interspersed repeats (SINE/LINE/LTR). RFhap strongly suppressed these errors, consistent with improved read-haplotype assignment, preventing phase switch across repeat-dense regions where markers are sparse or ambiguous. After these reductions, the residual long-switch errors occurred in tandem repeats/satellites and segmental duplications (SDs). Satellites present high repetitiveness and low unique marker density, while SDs are often high identical[25], making haplotype discrimination intrinsically difficult, particularly with lower per-read accuracy data (R9.4.1). The observed increase of SD-overlapping long-switch intervals in R9.4.1 assemblies likely reflects this fundamental limitation rather than a failure specific to RFhap; when paralogous copies are nearly identical, errors and missing markers can dominate the multi-k signal, and both graph decoration and read partitioning lose phasing precision.

From a user perspective, RFhap is a useful tool that improves phasing using standard trio data and an alignment-free k-mer lookup, with modest compute and memory requirements and a reproducible Nextflow implementation. This lowers the barrier to more accurate diploid assemblies without requiring additional assays (Hi-C, Strand-seq) or ultra-high coverage. Still, the residual phasing errors suggest clear next steps. First, SDs and satellite regions likely require either higher per-read accuracy, orthogonal long-range linkage (Hi-C/Strand-seq), or assembly-graph–aware integration of RFhap labels (rather than pure pre-assembly partitioning). Second, consensus accuracy under high-quality chemistries may benefit from post-assembly polishing that leverages both partitioned and pooled reads while maintaining haplotype constraints. Finally, extending the classifier with features tailored to duplication-rich contexts (e.g., copy-specific k-mers, local uniqueness scores, or repeat-aware priors) may help reduce the remaining burden of long-switch errors in the most challenging genomic regions.

In summary, our results demonstrate that RFhap is a practical contribution toward automated diploid telomere-to-telomere assemblies of human genomes.

## Methods

### Datasets

All datasets were obtained from the Human Pangenome Reference project[13] (Supplementary Table 1). We assembled four trios (HG002, HG005, HG01109, and HG01243), downloading parental short-read data and child Oxford Nanopore (ONT) long-read data. Trio relationships were validated with NGScheckMate[26], and basic read statistics were computed with seqkit[27].

### RFhap algorithm

RFhap is composed of five core modules: (i) identification of parental-specific k-mers, (ii) k-mer lookup in long reads, (iii) random-forest training and haplotype prediction, (iv) haplotype-specific read partitioning, and (v) haplotype de novo assembly.

### Identification of parental-specific k-mers from short-read data

Parental-specific *k*-mers are identified from parental short-read data using the k-mer counter KMC3[21]. For each parent, we built a KMC database of all *k*-mers (at k-mer lengths: 24, 27, 30, and 33). To filter sequencing errors and repetitive *k*-mers, we compute a *k*-mer count histogram using kmc_tools[21] and automatically select a parent-specific count range (c_min, c_max) by taking (i) the first local minimum after the low-frequency error peak as (c_min) and (ii) the first coverage where the histogram fell below that minimum count as (c_max). Parent-exclusive *k*-mers are then obtained by database subtraction using kmc_tools and dumped to text using kmc_dump[21], restricted to the parent-specific frequency interval (c_min, c_max), yielding the maternal-only and paternal-only *k*-mer sets used for downstream long-read classification.

### K-mer lookup in child long reads

To scan parental-specific k-mers along child long reads, we implemented FASTKM, a C++ program that performs high-throughput *k*-mer membership queries against one or more parental-specific *k*-mer sets. FASTKM takes as input (i) a configuration file listing the *k*-mer set file, its *k*-mer size, and the associated haplotype label, (ii) a gzipped long-read FASTQ/FASTA file, and (iii) the number of CPUs. For each *k*-mer set, FASTKM reads the list of parental-specific *k*-mers and converts each *k*-mer into a 64-bit integer representation using ntHash[22]. These hashed keys are used to construct a minimal perfect hash function (MPHF) with BooPHF[23], enabling constant-time mapping from a hashed *k*-mer to a unique array index. To avoid false positives of MPHF queries on non-members, we additionally store a compact fingerprint per key in a probabilistic set[24] (16-bit fingerprint; expected false-positive rate ≈ 1/2^16 ∼ 0.00001525878906). Each *k*-mer set is therefore represented by: (i) an MPHF over hashed keys and (ii) a fingerprint table used to validate membership after MPHF lookup. FASTKM streams long reads from the fastq/fasta long-read file using the kseq library[28]. Reads shorter than 500 bp are discarded. For each retained read and for each indexed *k*-mer set, FASTKM scans all *k*-mer of the read using ntHash to compute forward and reverse-complement hash values in O(1) per sliding window. At each position, FASTKM queries the MPHF with the forward hash and validates membership with the fingerprint table; if not found, it repeats the query using the reverse-complement hash. Positions with successful membership validation are recorded as marker hits. For each read × *k*-mer-set pair, FASTKM reports: (i) the number of matched markers (*n*), (ii) the mean (*m*) and standard deviation (*s*) of inter-marker distances (computed from consecutive hit positions), and (iii) an approximate marker span coverage (*cov*), defined as (lastHit-firstHit)/readLen×100. These features are stored as a tab-delimited matrix, with one row per read. Each row begins with the read identifier and read length, followed by the statistics for each *k-mer-haplotype* database. Finally, FASTKM parallelizes parental-specific scanning using a producer–consumer design with pthreads. A single producer thread streams reads into an in-memory buffer (∼ 5,000 reads), while multiple consumer threads (equal to the requested CPU count) dequeue batches and compute per-read statistics. Output writing is synchronized with a mutex to preserve correct formatting.

### RandomForest for haplotype read classification

The FASTKM per-read feature matrix is used to train and apply a multiclass random forest classifier [29] that assigns each long read to haplotype A, haplotype B, or unknown (U). From FASTKM values, we compute the total marker support per haplotype across k and A/B ratios using a pseudocount (optionally scaled by the number of available markers per haplotype). For training, reads with low overall marker support (total A + total B < 5) were labeled U. Otherwise, provisional A/B/U classes are assigned independently for each k and for the total ratio, and the consensus label per read is defined by majority vote across these five provisional classes (ties are assigned as U). We train on a random subset of at most 150,000 reads. Training is done on a balanced subset split into 75% training and 25% test sets, and model metrics are obtained using caret on a confusion matrix. The trained model, confusion matrix, and feature importance are stored for later use. Finally, the saved model is applied to all reads in the dataset to obtain per-read class labels and class probabilities (type=“prob”). Output tables report read ID, read length, predicted class, predicted probabilities, and the features used by the classifier.

### Nextflow pipeline of RFhap

RFhap is implemented as a Nextflow[30] pipeline (Figure S1) that executes all modules in a containerized and reproducible manner. Inputs are paternal and maternal short-read libraries and child long reads (provided as CSV lists), k-mer sizes (default: 24, 27, 30, 33), and an output directory. The workflow (i) builds parental-specific k-mer sets per k with KMC, (ii) formats and sorts the k-mer database descriptor, (iii) runs FASTKM to compute the per-read feature matrix, (iv) trains a random-forest model on a random subset of up to 150,000 reads, (v) predicts haplotype labels in parallel over the full dataset by splitting the matrix into chunks, (vi) converts predictions into read-ID lists for haplotype A/B/U, and (vii) extracts haplotype-partitioned FASTQ files with seqtk. Pipeline resources and execution are defined in the Nextflow configuration. Default parameters include num_cases=150000 for model training and size=200000 reads per prediction chunk. Process-level resource requests are configured on a per-step basis. The pipeline automatically produces standard Nextflow outputs (timeline, report, trace, and DAG) under the pipeline_info subdirectory.

### Assembly of read partitions

RFhap haplotype partitions were assembled separately with hifiasm v0.25.0 in --ont mode (haplotype A/maternal and haplotype B/paternal), adding the U (unknown) reads to each partition. As a baseline, we performed trio-binning directly with hifiasm v0.25.0 in --ont mode. Parental k-mer databases were first built using yak v0.1[21] with the yak count command (parameters: -b37 -t16) on the paternal and maternal paired-end short reads, producing pat.yak and mat.yak. These databases were then provided to hifiasm via -1 pat.yak and -2 mat.yak to generate a trio-aware assembly from the unpartitioned child long reads (hifiasm --ont -t32 -o <prefix> -1 pat.yak -2 mat.yak <child_reads>). Assembly metrics were computed with gfastats[31].

### Benchmark of haplotype-aware assemblies

To benchmark haplotype accuracy and contiguity, we evaluated all assemblies using Merqury [20]. Short-read k-mer databases for the mother, father, and child were built with meryl v1.3[20], and haplotype-specific markers (hap-mers) were derived from the parental sets. Using these hap-mers, Merqury computed haplotype-aware metrics for each assembly, including switch error (rate of haplotype error), phased NG50/N50 (computed over phase-consistent blocks), and phase-block statistics (e.g., average block size) as indicators of haplotype-resolved continuity and quality. In Merqury, switch events are stratified into short (transient, single-marker flips that revert within 200 bp) versus long (sustained flips spanning multiple consecutive markers and/or extending >200 bp), using shortNum=1 and shortLimit=200 in MerToPhaseBlock.java code.

The same evaluation was applied to RFhap-derived assemblies and the trio-mode hifiasm baseline. The benchmark was automated with a dedicated Nextflow pipeline (Figure S7). The workflow validates all required inputs at launch (short reads for both parents and the child, child long reads, and metadata such as method and dataset), and supports two modes: (i) from_assembly, where precomputed haplotype assemblies (FASTA) and assembly graphs (GFA without sequence) are provided; or (ii) from reads, where haplotype read-ID lists (A/B/U) are used to assemble haplotypes from the child long reads (assemble module). For Merqury, the pipeline optionally builds k-mer databases from parental/child short reads with meryl, derives hap-mers, and computes haplotype-aware assembly metrics. In all cases, basic contiguity metrics (e.g., N50) are extracted from the no-sequence GFA files using gfastats. Finally, results from all enabled evaluators are merged into a single summary table annotated with the specified method and dataset.

## Data availability

All sequencing datasets for trios HG002, HG005, HG01109, and HG01243 are listed in Supplementary Table 1, including download sources/links and basic read metrics. The hifiasm and RFhap assemblies have been deposited in Zenodo (DOI: https://doi.org/10.5281/zenodo.18261083).

## Code availability

RFhap source code is freely available at https://github.com/digenoma-lab/rfhap. The benchmarking workflow is available at https://github.com/digenoma-lab/Eval-RF-hap. Scripts used for figure generation and analyses are available at https://github.com/digenoma-lab/rfhap_ms.

## Author Contributions

Conceptualization: A.D.G.; Methodology: D.G., G.C., C.M., F.S., A.D.G.; Software: D.G., G.C., A.D.G.; Validation: D.G., G.C., J.F.M., A.D.G.; Formal analysis: D.G., A.D.G.; Investigation: D.G., G.C., J.F.M., C.M., F.S.; Resources: A.D.G.; Data curation: D.G., G.C.; Writing – Original Draft: D.G., A.D.G.; Writing – Review & Editing: D.G., G.C., J.F.M., C.M., F.S., A.D.G.; Visualization: D.G., A.D.G.; Supervision: F.S., A.D.G.; Project administration: A.D.G.; Funding acquisition: C.M., A.D.G. All authors have read and agreed to the published version of the manuscript.

## Funding

A.D.G. Fondecyt Nº 1221029. C.M. Fondecyt Iniciación Nº 11251927. Powered@NLHPC: This research was partially supported by the supercomputing infrastructure of the NLHPC (ECM-02) and the supercomputing infrastructure of the High-Performance Computing UOH laboratory (FIC 40059065-0) of the University of O’Higgins. A.D.G., C.M., F.S. and C.G. Centro UOH de Bioingeniería.

## Notes

https://doi.org/10.5281/zenodo.18261083

### Competing Interest Statement

The authors have declared no competing interest.

## References

1. Nurk S, Koren S, Rhie A, Rautiainen M, Bzikadze AV, Mikheenko A, et al. The complete sequence of a human genome. Science. 2022;376: 44–53.

2. Wang Y, Zhao Y, Bollas A, Wang Y, Au KF. Nanopore sequencing technology, bioinformatics and applications. Nat Biotechnol. 2021;39: 1348–1365.

3. Wenger AM, Peluso P, Rowell WJ, Chang P-C, Hall RJ, Concepcion GT, et al. Accurate circular consensus long-read sequencing improves variant detection and assembly of a human genome. Nat Biotechnol. 2019;37: 1155–1162.

4. Logsdon GA, Ebert P, Audano PA, Loftus M, Porubsky D, Ebler J, et al. Complex genetic variation in nearly complete human genomes. Nature. 2025;644: 430–441.

5. Cheng H, Concepcion GT, Feng X, Zhang H, Li H. Haplotype-resolved de novo assembly using phased assembly graphs with hifiasm. Nat Methods. 2021;18: 170–175.

6. Rautiainen M, Nurk S, Walenz BP, Logsdon GA, Porubsky D, Rhie A, et al. Telomere-to-telomere assembly of diploid chromosomes with Verkko. Nat Biotechnol. 2023;41: 1474–1482.

7. Mastrorosa FK, Miller DE, Eichler EE. Applications of long-read sequencing to Mendelian genetics. Genome Med. 2023;15: 42.

8. Sakamoto Y, Sereewattanawoot S, Suzuki A. A new era of long-read sequencing for cancer genomics. J Hum Genet. 2020;65: 3–10.

9. Kronenberg ZN, Rhie A, Koren S, Concepcion GT, Peluso P, Munson KM, et al. Extended haplotype-phasing of long-read de novo genome assemblies using Hi-C. Nat Commun. 2021;12: 1935.

10. Cheng H, Jarvis ED, Fedrigo O, Koepfli K-P, Urban L, Gemmell NJ, et al. Haplotype-resolved assembly of diploid genomes without parental data. Nat Biotechnol. 2022;40: 1332–1335.

11. Hansen NF, Dwarshuis N, Ji HJ, Rhie A, Loucks H, Logsdon GA, et al. A complete diploid human genome benchmark for personalized genomics. bioRxivorg. 2025. doi:10.1101/2025.09.21.677443

12. Zook JM, Catoe D, McDaniel J, Vang L, Spies N, Sidow A, et al. Extensive sequencing of seven human genomes to characterize benchmark reference materials. Sci Data. 2016;3: 160025.

13. Wang T, Antonacci-Fulton L, Howe K, Lawson HA, Lucas JK, Phillippy AM, et al. The Human Pangenome Project: a global resource to map genomic diversity. Nature. 2022;604: 437–446.

14. Sanders AD, Falconer E, Hills M, Spierings DCJ, Lansdorp PM. Single-cell template strand sequencing by Strand-seq enables the characterization of individual homologs. Nat Protoc. 2017;12: 1151–1176.

15. Belton J-M, McCord RP, Gibcus JH, Naumova N, Zhan Y, Dekker J. Hi–C: A comprehensive technique to capture the conformation of genomes. Methods. 2012;58: 268–276.

16. Rautiainen M. Ribotin: automated assembly and phasing of rDNA morphs. Bioinformatics. 2024;40. doi:10.1093/bioinformatics/btae124

17. Mastoras M, Asri M, Brambrink L, Hebbar P, Kolesnikov A, Cook DE, et al. Highly accurate assembly polishing with DeepPolisher. Genome Res. 2025;35: 1595–1608.

18. Hu J, Wang Z, Liang F, Liu S-L, Ye K, Wang D-P. NextPolish2: A repeat-aware polishing tool for genomes assembled using HiFi long reads. Genomics Proteomics Bioinformatics. 2024;22. doi:10.1093/gpbjnl/qzad009

19. Koren S, Rhie A, Walenz BP, Dilthey AT, Bickhart DM, Kingan SB, et al. De novo assembly of haplotype-resolved genomes with trio binning. Nat Biotechnol. 2018;36: 1174–1182.

20. Rhie A, Walenz BP, Koren S, Phillippy AM. Merqury: reference-free quality, completeness, and phasing assessment for genome assemblies. Genome Biol. 2020;21: 245.

21. Kokot M, Dlugosz M, Deorowicz S. KMC 3: counting and manipulating k-mer statistics. Bioinformatics. 2017;33: 2759–2761.

22. Kazemi P, Wong J, Nikolić V, Mohamadi H, Warren RL, Birol I. ntHash2: recursive spaced seed hashing for nucleotide sequences. Bioinformatics. 2022;38: 4812–4813.

23. Limasset A, Rizk G, Chikhi R, Peterlongo P. Fast and scalable minimal perfect hashing for massive key sets. arXiv [cs.DS]. 2017. doi:10.48550/ARXIV.1702.03154

24. Marchet C, Lecompte L, Limasset A, Bittner L, Peterlongo P. A resource-frugal probabilistic dictionary and applications in bioinformatics. arXiv [cs.DS]. 2017. doi:10.48550/ARXIV.1703.00667

25. Vollger MR, Dishuck PC, Sorensen M, Welch AE, Dang V, Dougherty ML, et al. Long-read sequence and assembly of segmental duplications. Nat Methods. 2019;16: 88–94.

26. Lee S, Lee S, Ouellette S, Park W-Y, Lee EA, Park PJ. NGSCheckMate: software for validating sample identity in next-generation sequencing studies within and across data types. Nucleic Acids Res. 2017;45: e103.

27. Shen W, Le S, Li Y, Hu F. SeqKit: A cross-platform and ultrafast toolkit for FASTA/Q file manipulation. PLoS One. 2016;11: e0163962.

28. Li H. seqtk: Toolkit for processing sequences in FASTA/Q formats. Github; Available: https://github.com/lh3/seqtk

29. Liaw A, Wiener M. Classification and Regression by randomForest. [cited 21 Jan 2025]. Available: https://journal.r-project.org/articles/RN-2002-022/RN-2002-022.pdf

30. Di Tommaso P, Chatzou M, Floden EW, Barja PP, Palumbo E, Notredame C. Nextflow enables reproducible computational workflows. Nat Biotechnol. 2017;35: 316–319.

31. Formenti G, Abueg L, Brajuka A, Brajuka N, Gallardo-Alba C, Giani A, et al. Gfastats: conversion, evaluation and manipulation of genome sequences using assembly graphs. Bioinformatics. 2022;38: 4214–4216.

