## Supplementary material for "Accurate haplotype-resolved *de novo* assembly of human genomes with RFhap"

†Corresponding author.

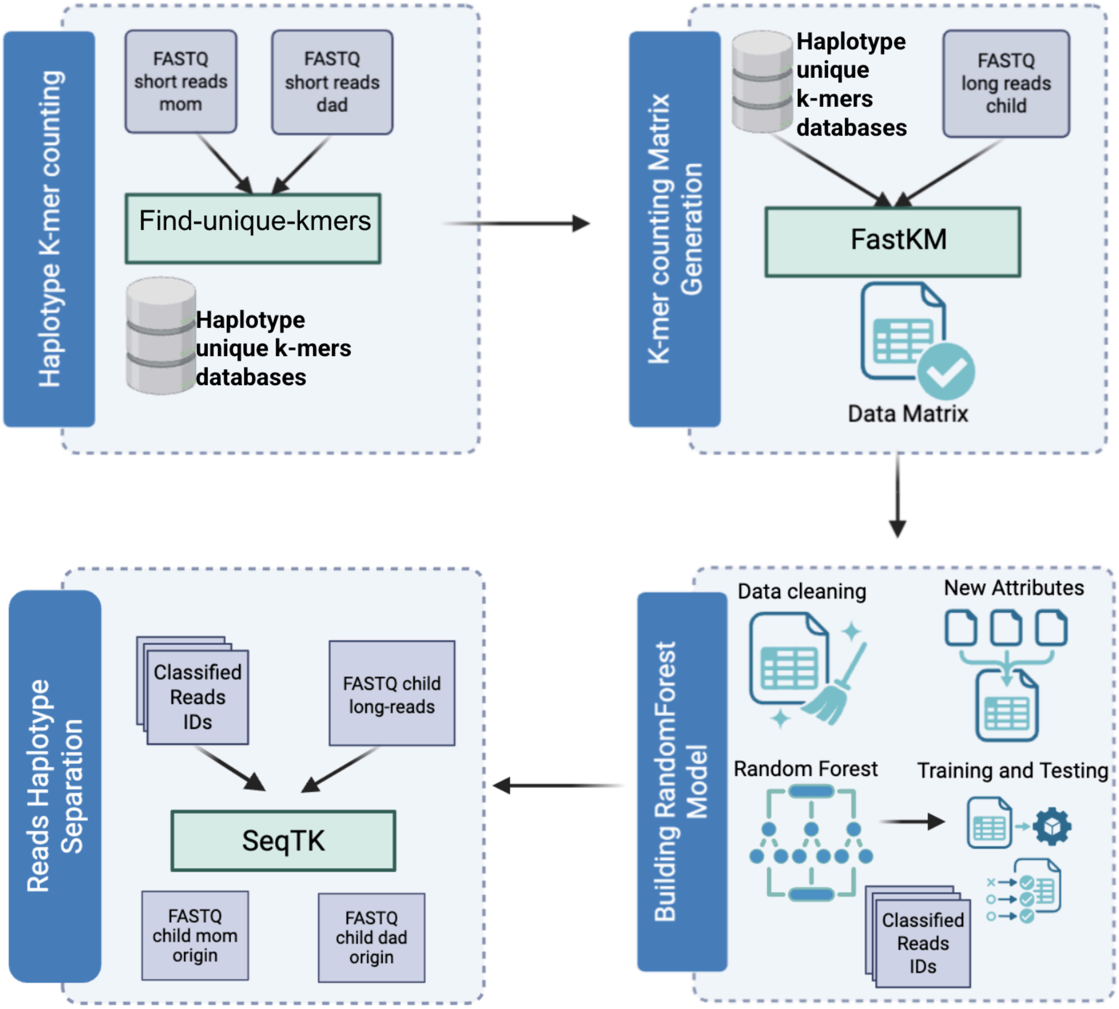


**Supplementary Figure 1: RFhap nextflow pipeline.** Parental short reads (FASTQ) are used to identify haplotype-specific k-mers, generating haplotype-unique k-mer databases. These databases, together with the child's long reads, are used by FastKM to compute a k-mer count matrix. After importing the data and deriving features, a Random Forest model is trained and applied to classify child long reads by haplotype. Classified read identifiers are then used by SeqTK to partition the child's long reads into maternal- and paternal-origin FASTQ files

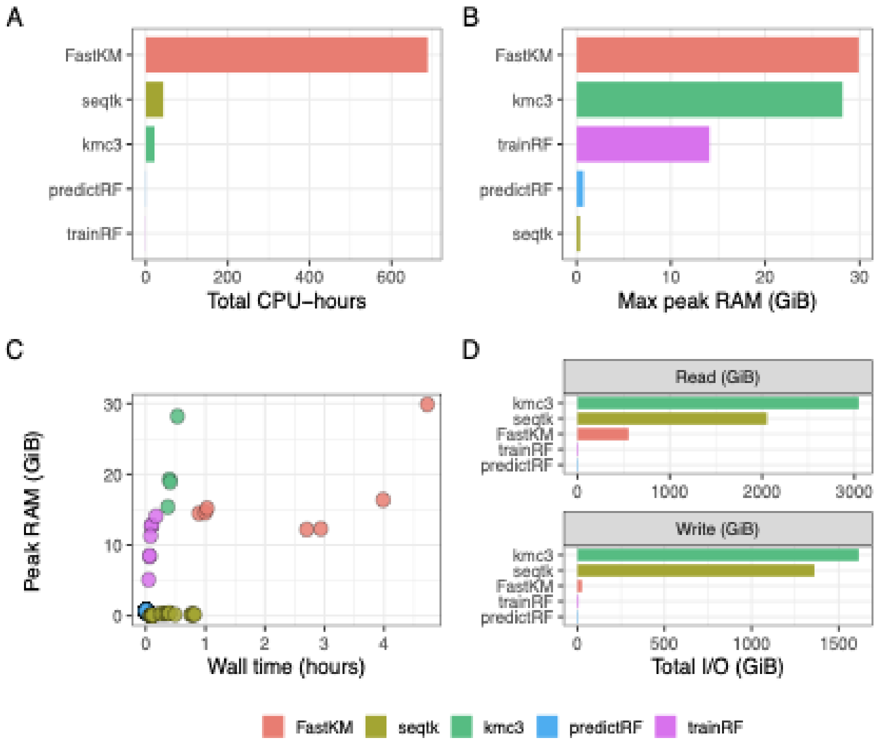


**Supplementary Figure 2: Computational requirements of the RFhap pipeline.**
(A) Total CPU-hours per process. (B) Maximum peak RAM (GiB) per process. (C) Task-level wall time vs peak RAM. (D) Total read/write I/O (GiB) per process. Only processes with ≥1 CPU-hour are shown. Computational Resources for binning the HG002 dataset.

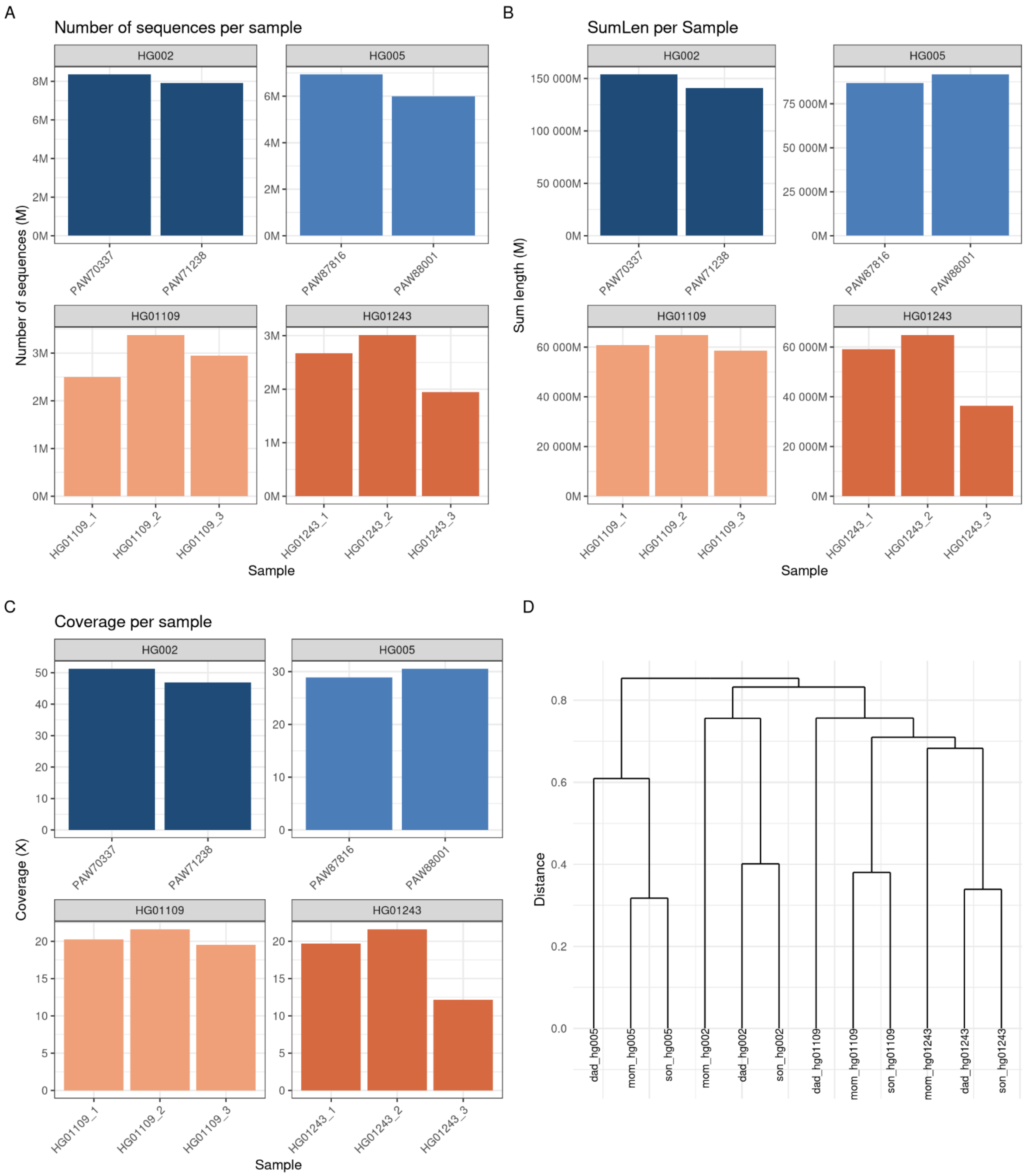


**Supplementary Figure 3: Sequencing summary statistic and sample concordance across datasets.** (A) Number of long-read sequences per sample for each dataset (HG002, HG005, HG01109, and HG01243). (B) Total sequenced length (SumLen) per sample, expressed in millions of bases(Mb). (C) Estimated long-read sequencing coverage per sample. and (D) NGScheckMate dendrogram showing concordance of related samples.

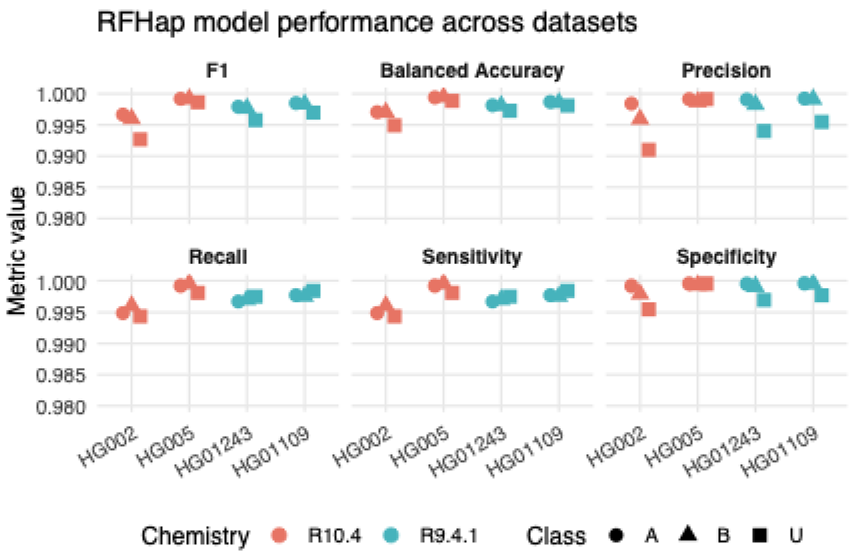


**Supplementary Figure 4. Performance of the RFhap classification model across sequencing datasets and chemistries.** Panels report F1 score, balanced accuracy, precision, recall, sensitivity, and specificity for four human datasets. Points are stratified by sequencing chemistry (R10.4 in red and R9.4.1 in blue) and by class label (A, B, and U; indicated by different point shapes).

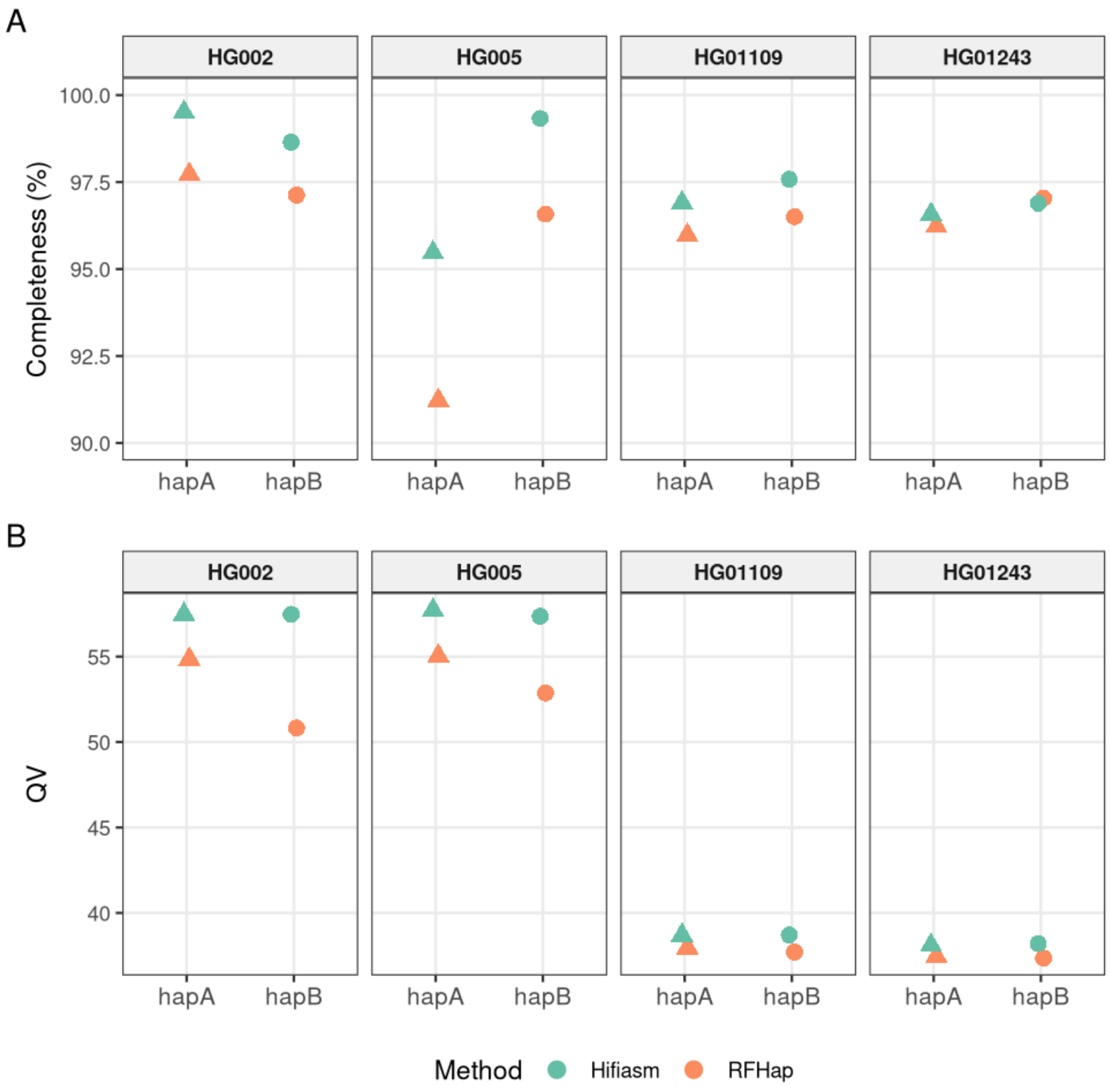


**Supplementary Figure 5.** Completeness and consensus quality assessment of haplotype-resolved assemblies generated with Hifiasm and RFhap.(A) Completeness (%) for each haplotype (hapA and hapB) across the HG002, HG005, HG01109, and HG01243 datasets. Triangles and circles indicate evaluation using maternal and paternal hapmer sets, respectively. Colors represent the assembly method: Hifiasm (pink) and RFhap (turquoise). (B) Merqury Quality Value (QV) for the same assemblies, reporting the Phred-scaled base error rate, where higher values indicate higher consensus accuracy.

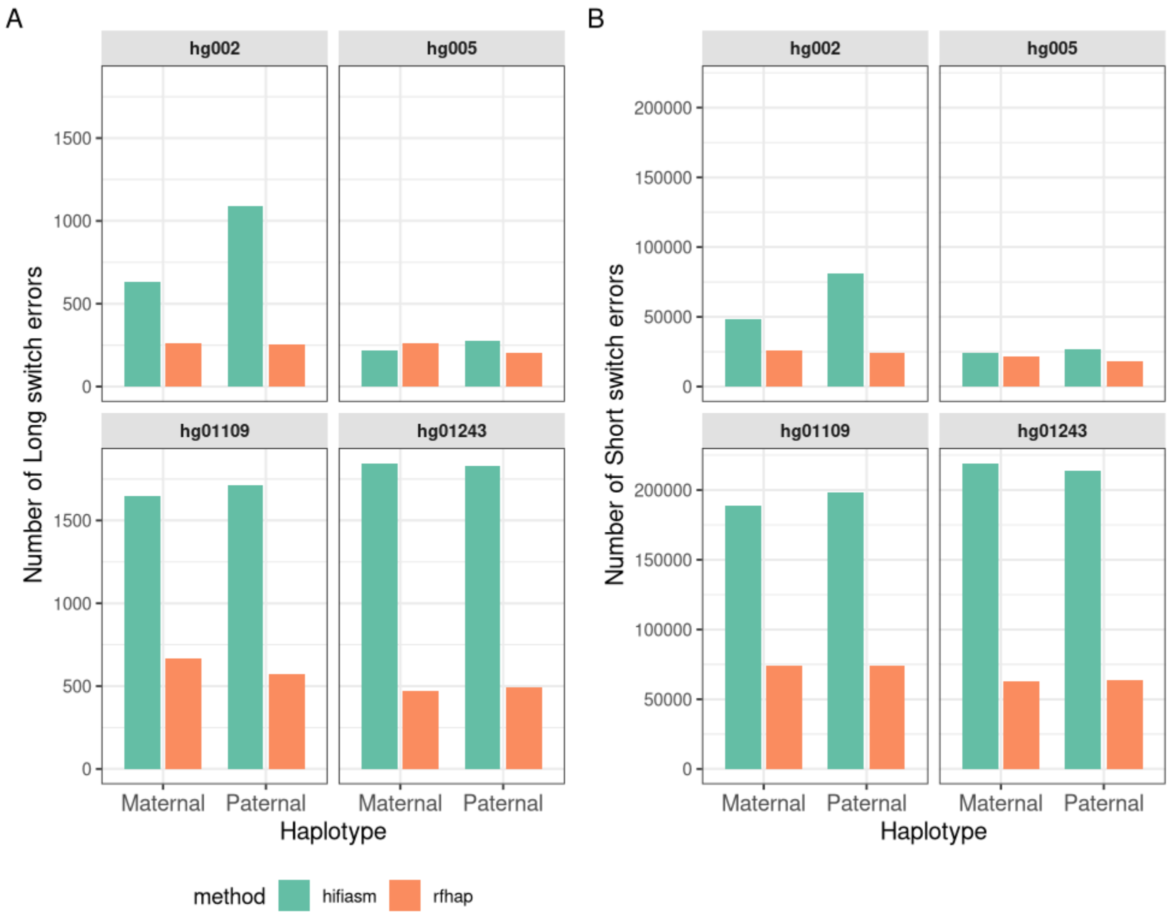


**Supplementary Figure 6. Comparison of switch error counts across haplotypes, datasets, and phasing methods.** (A) Number of long switch errors and (B) number of short switch errors, for maternal and paternal haplotypes across four human datasets. Bars are colored by phasing method (HiFiAsm in blue and RFhap in orange).

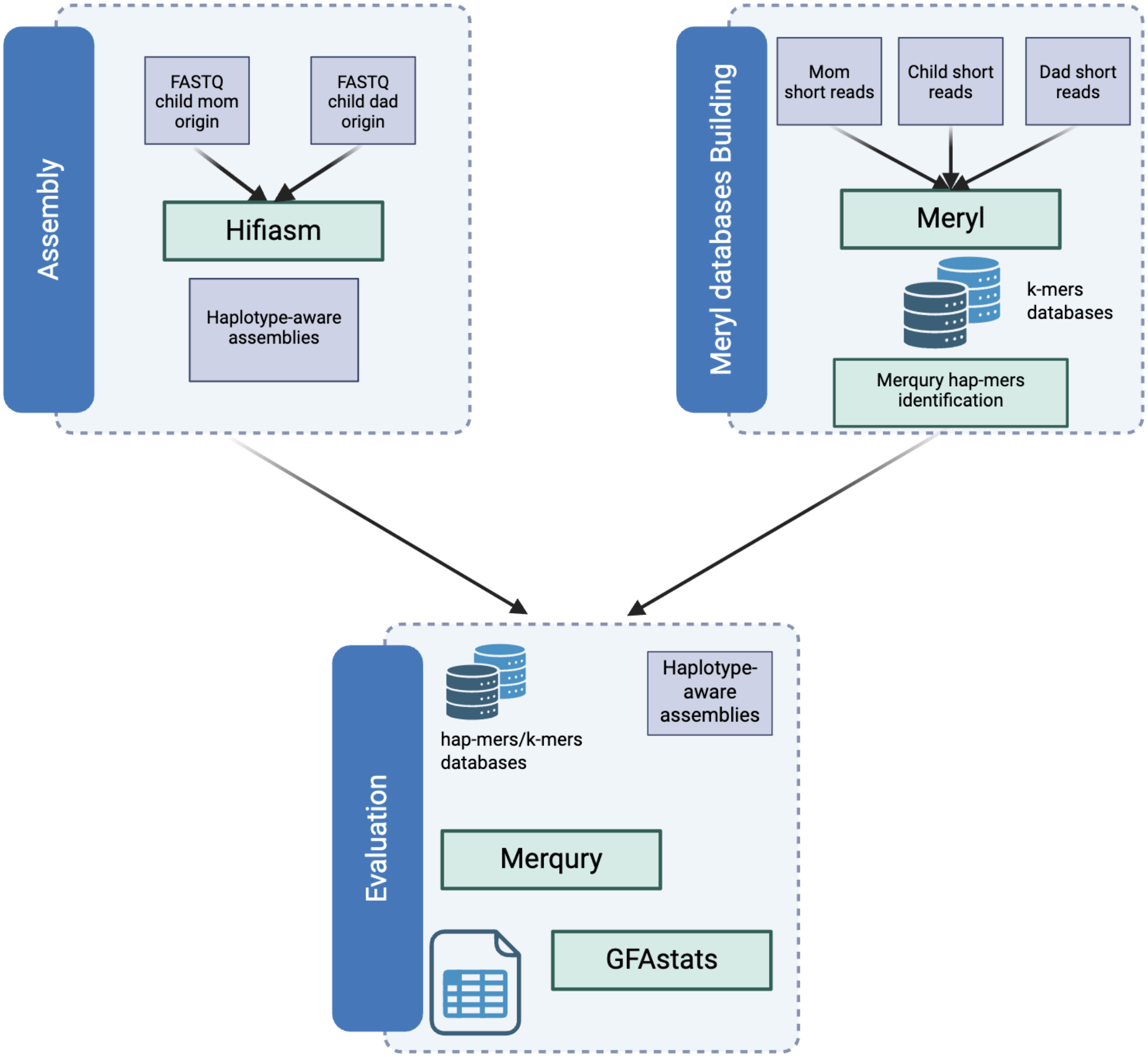


**Supplementary Figure 7: Benchmark nextflow pipeline.** The workflow processes parental and child short reads with Meryl to build hap-mer databases, then evaluates the resulting haplotype assemblies with Merqury, a k-mer–based toolkit that estimates assembly quality and completeness (QV and k-mer completeness) and quantifies haplotype phasing accuracy using hap-mers. GFAstats is used in parallel to summarize assembly statistics. The pipeline can also generate haplotype-aware assemblies from read partitions using hifiasm.

**Supplementary Table 1: Datasets for RFhap benchmark.** Trio identifiers, sequencing technology, and library (flow cell or paired-end), average read quality, read N50, and data source URL.

| Dataset | Trio | Technology | Flow-cell/pair-end | Avg. Quality | N50 (bp) | URL |
|---|---|---|---|---|---|---|
| HG01109 | maternal | Illumina | R1 | 24.7 | 151 | s3://human-pangenomics/NHGRI_UCSC_panel/HG01108/illumina/ |
| | | | R2 | 23.24 | | |
| | paternal | Illumina | R1 | 24.28 | 151 | s3://human-pangenomics/NHGRI_UCSC_panel/HG01107/illumina/ |
| | | | R2 | 23.45 | | |
| | child | Illumina | R1 | 24.01 | 150 | http://ftp.sra.ebi.ac.uk/vol1/fastq/ERR398/002/ERR3988842/ |
| | | | R2 | 26.01 | | |
| | child | ONT | HG01109_1_R941_Guppy_6.1.2 | 14.07 | 48,161 | s3://human-pangenomics/NHGRI_UCSC_panel/HG01109/nanopore/Guppy_6.1.2/ |
| | | | HG01109_2_R941_Guppy_6.1.2 | 14.41 | 52,856 | |
| | | | HG01109_3_R941_Guppy_6.1.2 | 14.2 | 48,326 | |
| HG002 | maternal | Illumina | R1 | 26.5 | 148 | https://s3-us-west-2.amazonaws.com/human-pangenomics/working/HPRC_PLUS/HG002/raw_data/Illumina/parents/HG004/ |
| | | | R2 | 21.27 | | |
| | paternal | Illumina | R1 | 25.67 | 148 | https://s3-us-west-2.amazonaws.com/human-pangenomics/working/HPRC_PLUS/HG002/raw_data/Illumina/parents/HG003/ |
| | | | R2 | 19.95 | | |
| | child | Illumina | R1 | 26.31 | 148 | https://s3-us-west-2.amazonaws.com/human-pangenomics/working/HPRC_PLUS/HG002/raw_data/Illumina/ |
| | | | R2 | 21.46 | | |
| | child l | ONT SUP | PAW70337 | 18.7 | 29395 | s3://ont-open-data/giab_2025.01/basecalling/sup/HG002/ |
| | | | PAW71238 | 18.93 | 29727 | |
| HG005 | maternal | Illumina | R1 | 25.37 | 148 | s3://human-pangenomics/working/HPRC_PLUS/HG005/raw_data/Illumina/parents/HG007/<br>Flowcells:<br><br>141008_D00360_0059_BHB66DADXX/<br>141015_D00360_0061_BHB66HADXX/ |
| | | | R2 | 19.75 | | |
| | paternal | Illumina | R1 | 25.53 | 148 | s3://human-pangenomics/working/HPRC_PLUS/HG005/raw_data/Illumina/parents/HG007/<br><br>Flowcells: 141008_D00360_0058_AHB675ADXX<br>141015_D00360_0060_AHB6D7ADXX |
| | | | R2 | 20.17 | | |
| | child | Illumina | R1 | 25.69 | 250 | s3://human-pangenomics/working/HPRC_PLUS/HG005/raw_data/Illumina/child// |
| | | | R2 | 22.69 | | |
| | child l | ONT SUP | PAW87816 | 19.03 | 23,909 | s3://ont-open-data/giab_2025.01/basecalling/sup/HG005/PAW87816/calls.sorted.bam |
| | | | PAW88001 | 19.7 | 25,174 | |
| HG01243 | maternal | Illumina | R1 | 24.28 | 151 | s3://human-pangenomics/NHGRI_UCSC_panel/HG01242/illumina/ |
| | | | R2 | 23.45 | | |
| | paternal | Illumina | R1 | 24.34 | 151 | s3://human-pangenomics/NHGRI_UCSC_panel/HG01241/illumina/ |
| | | | R2 | 23.46 | | |
| | child | Illumina | R1 | 24.49 | 150 | http://ftp.sra.ebi.ac.uk/vol1/fastq/ERR398/008/ERR3988858/ |
| | | | R2 | 22.46 | | |
| | child | ONT | HG01243_1_R941_Guppy_6.1.2 | 14.13 | 48,654 | s3://human-pangenomics/NHGRI_UCSC_panel/HG01243/nanopore/Guppy_6.1.2 |
| | | | HG01243_2_R941_Guppy_6.1.2 | 14.6 | 47,448 | |
| | | | HG01243_3_R941_Guppy_6.1.2 | 14.24 | 44,546 | |
